# Live-Cell Imaging of (p)ppGpp with RNA-based Fluorescent Sensors

**DOI:** 10.1101/2021.05.12.443921

**Authors:** Zhining Sun, Rigumula Wu, Bin Zhao, Rilee Zeinert, Peter Chien, Mingxu You

## Abstract

Guanosine tetra- and pentaphosphate, (p)ppGpp, are important alarmone nucleotides that regulate bacterial survival in stressful environment. A direct detection of (p)ppGpp in living cells is critical for our understanding of the mechanism of bacterial stringent response. However, it is still challenging to directly image and measure cellular (p)ppGpp. Here, we report a type of RNA-based fluorescent sensors for live-cell imaging of (p)ppGpp. Our sensor is engineered by conjugating a recently identified (p)ppGpp-specific riboswitch with a fluorogenic RNA aptamer, Broccoli. These sensors can be genetically encoded and enable direct monitoring of cellular (p)ppGpp accumulation. Unprecedented information on cell-to-cell variation and cellular dynamics of (p)ppGpp levels can now be obtained under different nutritional conditions. We predict that these RNA-based sensors will be broadly adapted to study bacterial stringent response.

---

(p)ppGpp, often referred to as the “Magic Spot”, is tightly involved in the stringent response in bacteria.^[1–3]^ When bacterial cells are under environmental threats, such as nutrient starvation, heat shock, and antibiotic treatment, (p)ppGpp is rapidly produced by RSH (RelA/SpoT homologs) enzymes and accumulates to high levels. As an important signaling molecule, (p)ppGpp allows the cells to redistribute resources towards amino acid synthesis and stress survival.^[4]^ (p)ppGpp can function by directly acting on RNA polymerases to alter their promoter selection and rate of transcription.^[5,6]^ It can also modulate the activity of a series of transcription factors, enzymes, and regulatory RNAs.^[7,8]^ Notably, (p)ppGpp was also recently identified to play an integral role in regulating biofilm formation, antibiotic resistance, and persistence.^[9,10]^

Even though (p)ppGpp has been shown to control an impressive array of pathways to reprogram bacteria during stress, one major gap is the lack of tools to directly detect (p)ppGpp in living cells. Traditional methods of detecting (p)ppGpp rely on radiolabeling (p)ppGpp with ^32^P and measuring with thin layer chromatography^[11–13]^ or high-performance liquid chromatography.^[14,15]^ Some colorimetric and fluorescent probes have also been developed.^[16–21]^ However, even though these chemosensors enable real-time monitoring of *in vitro* ppGpp synthesis using *E. coli* ribosomal complex, they are limited for solution-based detection. There is still a void in detecting ppGpp in live cells, especially at the single-cell level for studying cell-to-cell variations. Some fluorescent protein-based sensors have recently been developed attempting to image (p)ppGpp in live cells.^[22,23]^ This type of sensors does not directly target (p)ppGpp, but rather its activation system. Therefore, sensors that can directly measure (p)ppGpp levels in single cells is still highly demanded for understanding the bacterial stringent response and their adaption to antibiotics.

To fill this gap, we report here the development of genetically encodable RNA-based sensors for fluorescence imaging of (p)ppGpp in living cells. These RNA-based sensors have three major components: a (p)ppGpp-specific aptamer, a transducer sequence, and a fluorogenic RNA reporter, e.g., so-called “Broccoli”.^[24]^ Aptamers are single-stranded oligonucleotides that can bind their target with high affinity and selectivity. Naturally existing aptamers (i.e., riboswitches) that can selectively bind with (p)ppGpp have recently been discovered.^[25]^ In our sensor construct, when (p)ppGpp binds to the aptamer region, the transducer sequence hybridizes and facilitates the folding of the Broccoli RNA. The folded Broccoli can further bind to its cognate dye, DFHBI-1T, and activate its fluorescence (Figure 1a). As a result, the fluorescence signal can be used to detect (p)ppGpp.

**Figure 1.**
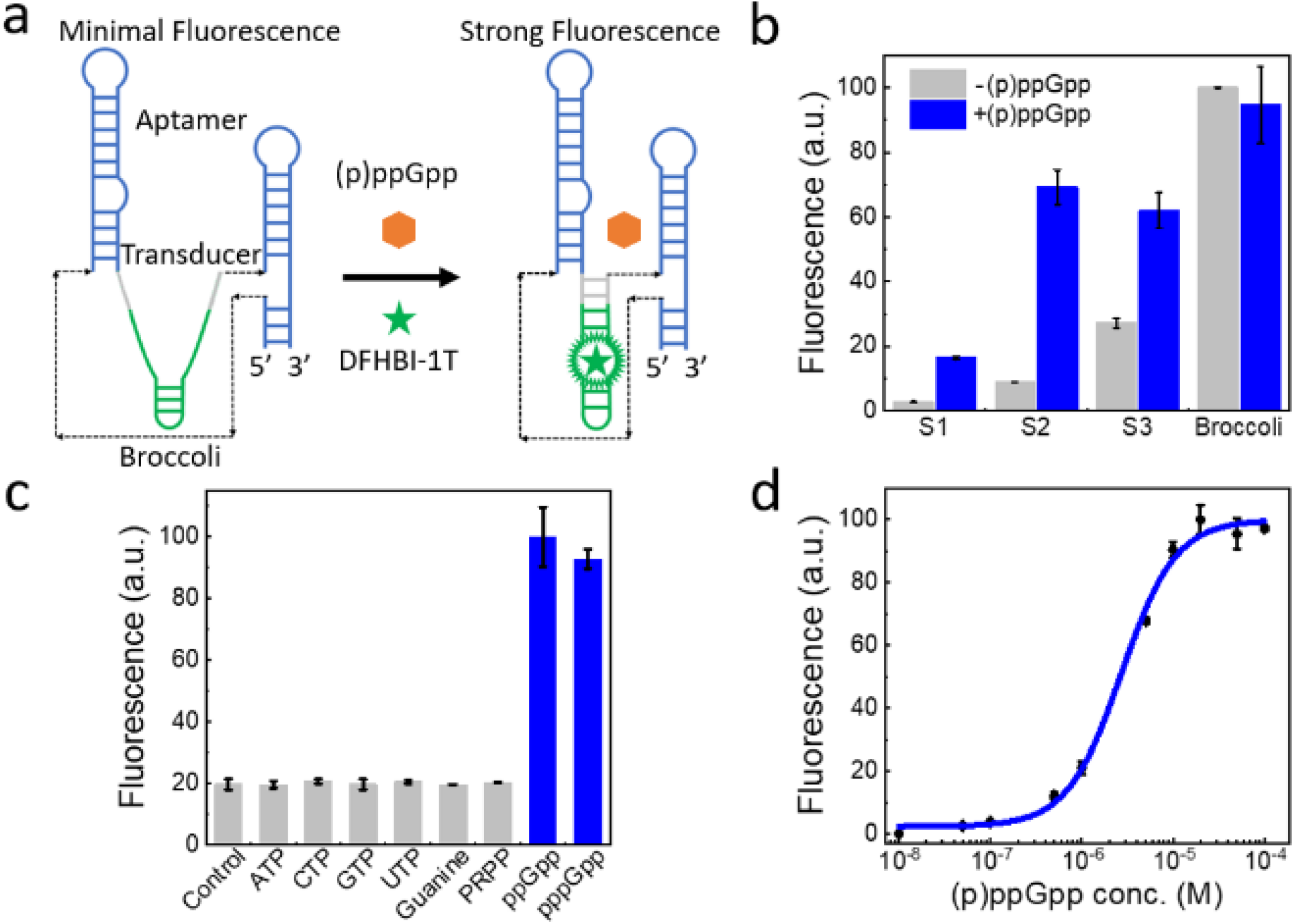
Design and *in vitro* characterization of the RNA-based (p)ppGpp sensor. **a)** Modular design of the sensors comprises of a Broccoli (green), a (p)ppGpp-binding aptamer (blue), and a transducer sequence (gray). **b)** Optimization of transducer sequences. The corresponding sequences were shown in Table S1. Spectra were measured in a solution containing 1 μM RNA, 20 μM DFHBI-1T, and either 0 or 10 μM (p)ppGpp, at 90 min after mixing. **c)** Selectivity of the S2 sensor. Spectra were measured with 10 mM NTPs or 10 μM other ligands in a solution containing 1 μM S2 and 20 μM DFHBI-1T. Control was measured without adding ligands. **d)** Dose-response curve of the S2 sensor. The limit of detection was calculated to be 0.3 µM. Shown are the mean ± standard deviation of three independent replicates.

We first designed three RNA sensor constructs by fusing Broccoli into the P0 stem region of the (p)ppGpp riboswitch through different transducer sequences. These transducers have been optimized based on an Mfold online software. However, minimal fluorescence activation was shown upon adding (p)ppGpp (Figure S1). After further examining the crystal structure of the riboswitch (PDB: 6DMC), we realized that the P2 stem region is very close to the (p)ppGpp-binding pocket and may be sequence-independent.^[26–28]^ To test if (p)ppGpp sensors could be developed by inserting Broccoli into the P2 stem, we designed another five transducer sequences based on the Mfold simulation (Table S1). Indeed, three of these constructs, S1–S3, exhibited respectively, 5.9-fold, 7.9-fold, and 2.3-fold fluorescence enhancement upon the addition of 10 µM ppGpp (Figure 1b and S2).

We chose S1 and S2 sensors for further *in vitro* tests due to their high fluorescence enhancement. We first tested their selectivity against several (p)ppGpp analogs (Figure 1c and S3). The S2 fluorescence can be similarly activated by either guanosine tetraphosphate (ppGpp) or guanosine pentaphosphate (pppGpp). Interestingly though, when using the S1 sensor, pppGpp activates 40% less fluorescence than ppGpp does (Figure S3). Both S1 and S2 exhibited no fluorescence activation in the presence of NTPs, guanine, or phosphoribosyl pyrophosphate (PRPP). S1 and S2 are indeed highly selective towards (p)ppGpp.

We then studied the detection range of S1 and S2. A dose-response curve was measured for each sensor after adding various concentrations of ppGpp (Figure 1d and S4). Our results indicated that S2 can effectively detect (p)ppGpp ranging from 0.5 μM to 10 μM, whereas S1, with a more stable transducer (2 bp vs. 1 bp) exhibits a broader detection range, 1–100 μM. These two sensors can generally cover the concentration range that (p)ppGpp is known to normally exhibit in bacterial cells.^[14,29]^ In order to potentially detect even higher concentrations of (p)ppGpp, we also designed several fluorogenic RNA sensors by adopting the aptamer domain from a *D. hafniense ilvE* riboswitch that has a weak binding towards (p)ppGpp. Indeed, an optimal sensor based on the new design, named D3, exhibited an even higher detection range as compared to the S-type sensors (Figure S4).

We have also determined the kinetics of RNA sensor activation and deactivation. In general, S2 exhibited a faster fluorescence activation than S1. After mixing ppGpp with S2, half-maximum fluorescence level was reached in 7 min, with 80% of maximal signal in 19 min (Figure S5). By contrast, it would take 65 min or 39 min for the S1 or D3 signal to reach half-maximum, and 205 min or 115 min for S1 or D3 to reach 80% of maximal fluorescence (Figure S6 and S7). S2 also exhibited a fast fluorescence deactivation rate, with >95% signal decrease occurs in less than 4 min after removing the free ppGpp (Figure S8). These results indicated that the fluorescence signal of the RNA-based sensors, especially the S2, can be used to monitor the dynamic variations of (p)ppGpp levels. Considering its relatively high fluorescence intensity and fast kinetic response, we continued with S2 for the subsequent studies.

To visualize (p)ppGpp in live cells, we first cloned S2 into a pET28c vector and transformed it into BL21 Star™ (DE3) *E. coli* cells. As controls, we also transformed the same type of cells with empty or Broccoli-expressing pET28c plasmids. Following incubation with DFHBI-1T, as expected, Broccoli-expressing cells showed high fluorescence signal, while cells expressing empty pET28c vector had minimum fluorescence (Figure 2a). Cells expressing the S2 sensor also exhibited low fluorescence signal, consistent with the expected low levels of (p)ppGpp in rich growth conditions. Importantly, upon induction of (p)ppGpp by a truncated RelA protein,^[4,30]^ strong fluorescence signal was observed in S2-expressing cells, which indicated that the S2 signal is indeed correlated with (p)ppGpp cellular concentrations (Figure 2a).

**Figure 2.**
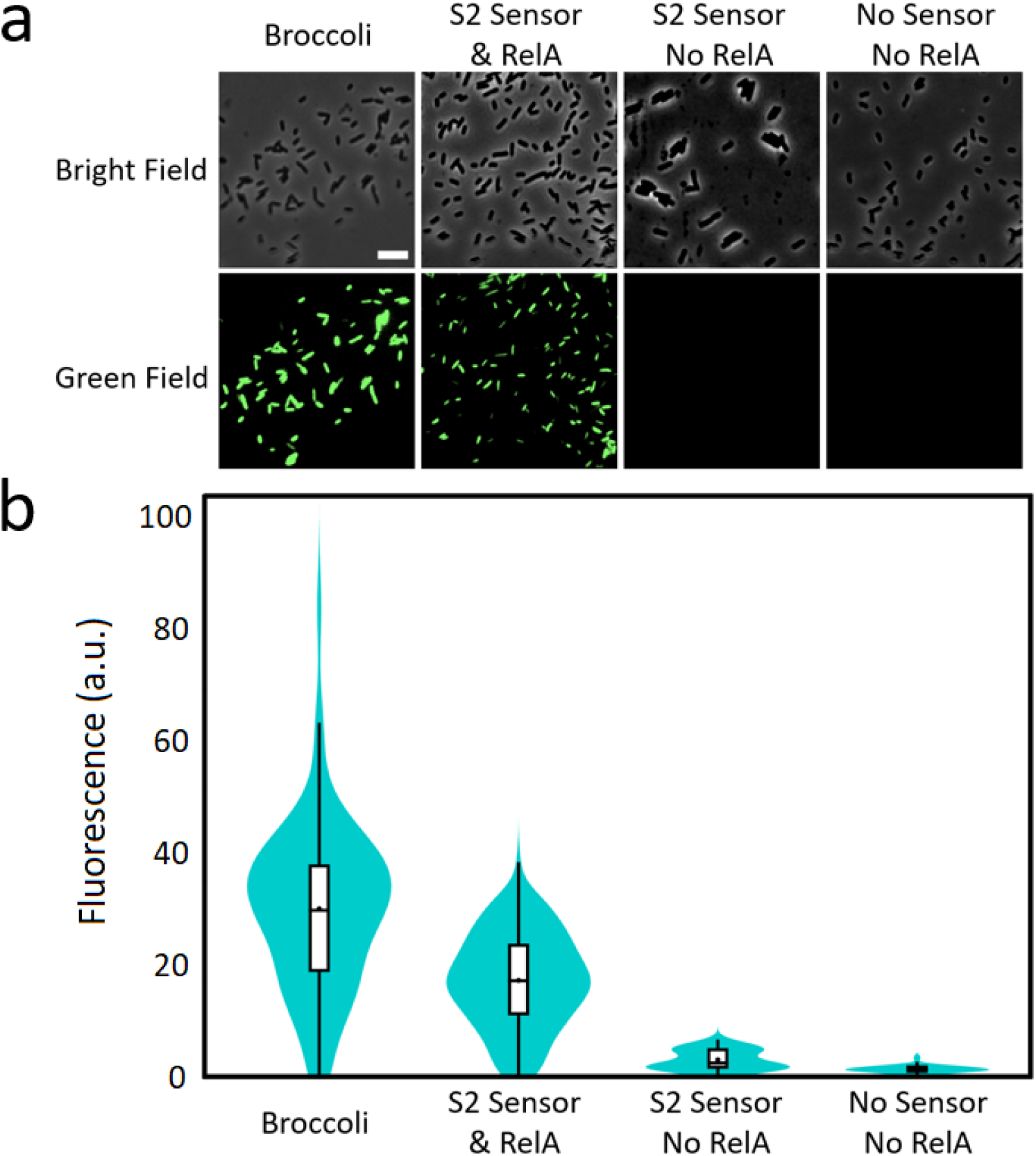
Live-cell imaging of (p)ppGpp with the S2 sensor. **a)** Confocal fluorescence imaging in live BL21 Star^™^ (DE3) cells. Images were taken 1 h after adding 200 µM DFHBI-1T. Both the cell morphology (bright field) and S2 fluorescence (green field) were shown. Scale bar, 10 μm. **b)** Distributions of the cellular fluorescence. In each condition, a total of >100 cells were measured from three experimental replicates. The width of violin represents the percentage of cells with the same fluorescence intensity. The white box region indicates the interquartile range, i.e., middle 50% of cellular fluorescence. The dot and short horizontal line in the box indicate the mean and median value of each data set.

We have further quantified the fluorescence intensities from hundreds of individual cells in each condition. Our results indicated that in strains with elevated (p)ppGpp, S2 fluorescence was approximately 70% of that of Broccoli (Figure 2b). A ∼7-fold fluorescence enhancement was observed if compared with that in the absence of truncated RelA expression. These data are highly consistent with that *in vitro* (Figure 1b).

To confirm if these RNA-based sensors can indeed detect intracellular (p)ppGpp, we conducted another series of tests under different nutritional stress. We first cultured S2-expressing *E. coli* cells in M9 minimal medium, where high cellular level of (p)ppGpp will be generated due to the low nutrient content of the media.^[31]^ As predicted, strong cellular fluorescence was observed (Figure 3a). After supplementing the M9 medium with nutrients casamino acids (CAA) and glucose, a dramatic and rapid fluorescence decrease was observed, indicating a decreased (p)ppGpp level in this nutrient-rich medium (Figure 3a). As a control, fluorescence signals of Broccoli-expressing cells did not change during the same nutrient supplementation process (Figure S9). We have further tested to first incubate S2-expressing cells in M9 medium containing both CAA and glucose, and then deprived the nutrients by switching to the M9 minimum medium. Again as expected, a significant increase in cellular fluorescence signal was observed (Figure 3a). All these data indicate that S2 can be indeed used to detect changes in the cellular (p)ppGpp level.

**Figure 3.**
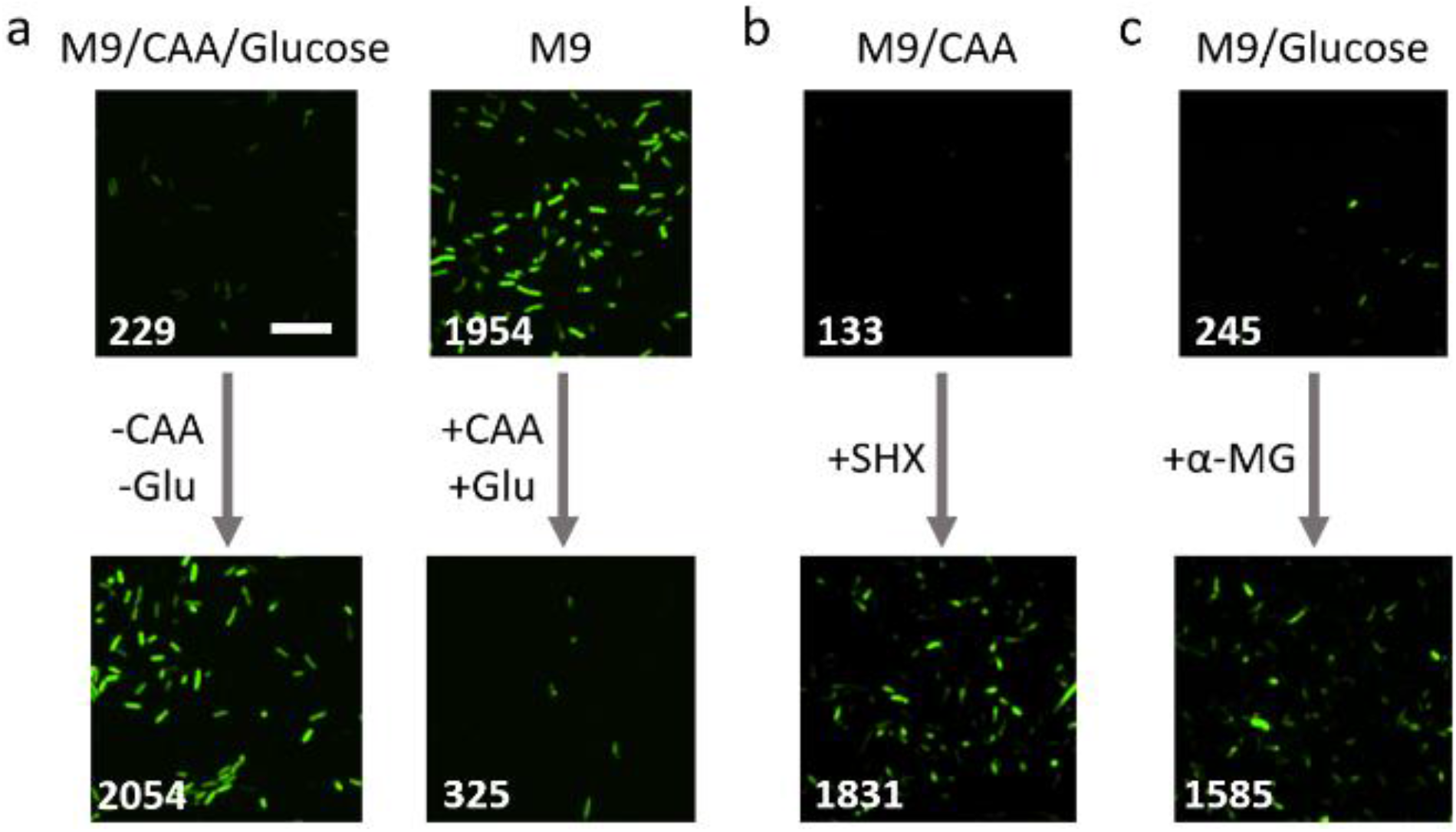
Imaging of (p)ppGpp biosynthesis under nutritional stress. **a)** Confocal fluorescence imaging of S2-expressing BL21 Star^™^ (DE3) cells in either M9 minimal medium or M9 medium supplemented with 0.2% casamino acids (CAA) and 0.4% glucose (Glu). Shown are images before or 15 min after changing the medium. Scale bar, 10 μm. **b)** Serine hydroxamate (SHX)-induced (p)ppGpp synthesis. These *E. coli* cells were first incubated in M9 medium supplemented with 0.2% CAA, and then 1% final concentration of SHX was added 15 min before imaging. **c)** Methyl-α-glucose (α-MG)-induced (p)ppGpp biosynthesis. These bacterial cells were first incubated in M9 medium supplemented with 0.4% glucose, and then 2.5% final concentration of α-MG was added 15 min before imaging. Numbers in each image indicated the averaged cellular fluorescence intensity.

We have further studied the effect of chemical inducers in the biosynthesis of (p)ppGpp. Serine hydroxamate (SHX) and methyl-α-glucose (α-MG) are two commonly used chemicals to trigger the intracellular accumulation of (p)ppGpp. SHX and α-MG functions by inducing the starvation of amino acids and glucose, respectively.^[32–34]^ Indeed, after adding 1% final concentration of SHX or 2.5% α-MG, significant increase in cellular S2 fluorescence signal was observed (Figure 3, S10, and S11). Compared to the effect of SHX, the addition of α-MG induced obviously less fluorescence enhancement. This result is consistent with some previous findings that the glucose starvation has less influence on the biosynthesis of (p)ppGpp than does the amino acids starvation.^[33,35]^

We also measured the kinetics of cellular (p)ppGpp accumulation in response to the medium exchange. S2-expressing cells were first grown in nutrient-rich medium and then switched into a nutrient-limited M9 minimum medium. By imaging every 10 min over 1 h, a time-dependent increase in the cellular fluorescence was observed. In most cells, a clear fluorescence activation can be seen within the first 20 min and then reached plateau (∼7-fold increase) at ∼60 min (Figure 4a and S12). To study if there is a cell-to-cell variation in (p)ppGpp biosynthesis during this medium switching process, we monitored the fluorescence signal change of over 100 individual cells (Figure 4b). On average, half-maximum fluorescence could be reached in 20 min. By defining fast kinetics as cellular fluorescence doubled within 20 min, the majority (84%) of cells exhibited fast (p)ppGpp accumulation rates, while 16% of cells showed a slow rate of (p)ppGpp biosynthesis. These data confirm that (p)ppGpp is rapidly accumulating in *E. coli* cells after encountering nutritional stress.

**Figure 4.**
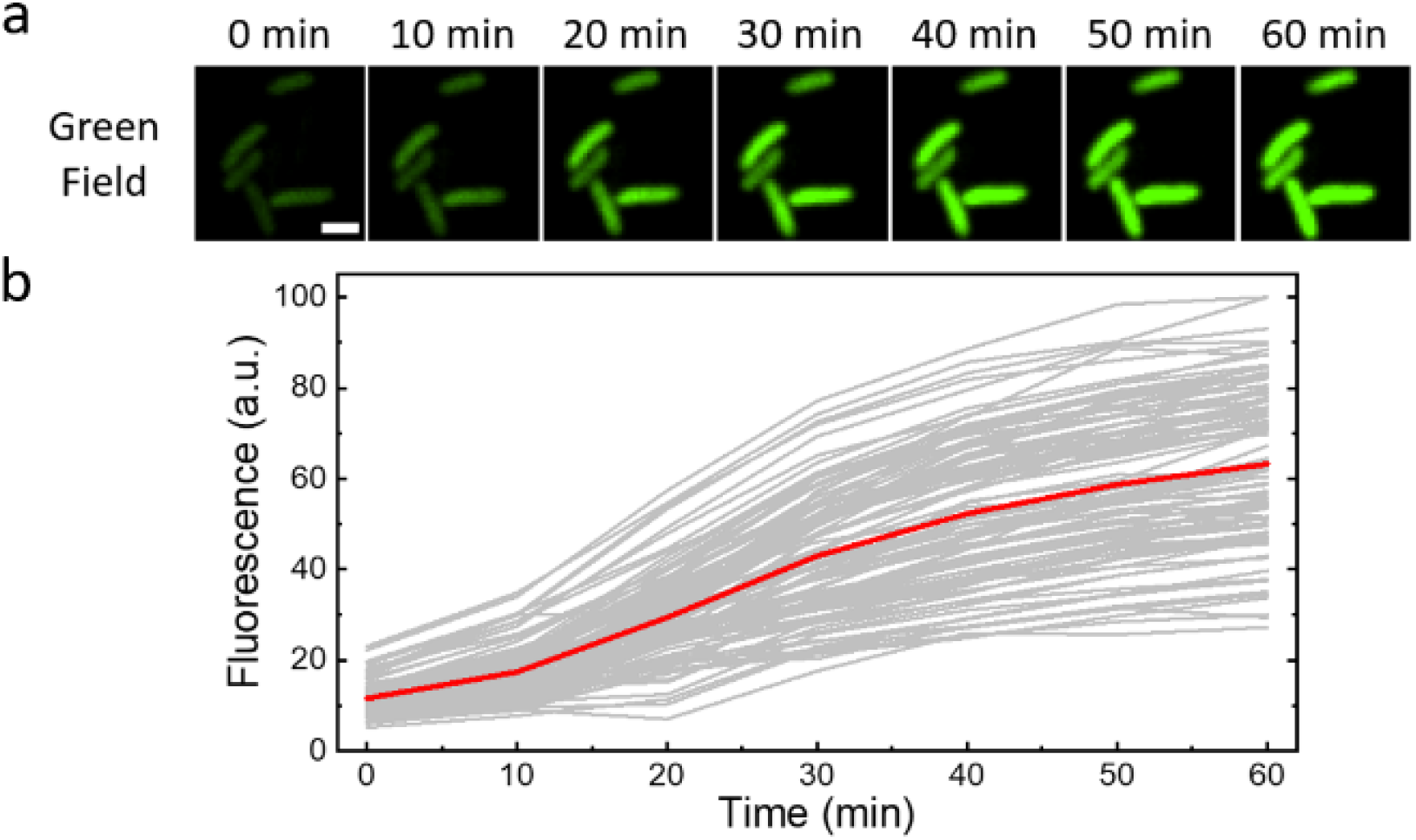
Kinetics of (p)ppGpp biosynthesis in *E. coli* cells. **a)** Representative confocal fluorescence images of S2-expressing BL21 Star^™^ (DE3) cells after switching at 0 min from M9/0.2% casamino acids/0.4% glucose medium to M9 minimal medium. **b)** Distribution of S2 cellular fluorescence levels as measured from 102 individual cells. Shown are the dynamic changes in the fluorescence signal of each individual cell (gray) and the averaged signal (red). Scale bar, 2 μm.

Lastly, we wanted to measure the cellular distributions of (p)ppGpp. We cloned the S2 sensor into a pETDuet vector that contains a separate promoter expressing eqFP670, a far-red fluorescent protein. Here, eqFP670 expression is used as a reference to normalize the variations of plasmid expression level among different cells. After transforming these plasmids into BL21 Star^™^ (DE3) cells, we incubated these cells in either M9/CAA/glucose medium or M9 minimal medium for 90 min. As expected, significantly higher S2 fluorescence exhibited in the nutrient-limited M9 minimal medium than that in M9/CAA/glucose medium (Figure 5a). There are some clear cell-to-cell variations in the plasmid expression level. We thus further measured S2 fluorescence at a single-cell level and normalized it to the corresponding eqFP670 fluorescence intensity (Figure 5b). In the nutrient-rich M9/CAA/glucose medium, >90% of cells exhibited minimum (p)ppGpp levels, i.e., Green/Red ratio (G/R) < 0.05, and only 2% of cells exhibited moderate (p)ppGpp biosynthesis (G/R > 0.10). By contrast, when nutrients are removed, cellular (p)ppGpp level distribution was largely symmetric. The majority (71%) of cells showed moderate (p)ppGpp levels (G/R, 0.05–0.25), while 11% and 18% of cells showed minimum (G/R < 0.05) and high (G/R > 0.25) (p)ppGpp levels, respectively. The highly accumulated (p)ppGpp in a small portion (∼1/5) of cells may be potentially interesting targets for the study of biofilm formation and persistence.

**Figure 5.**
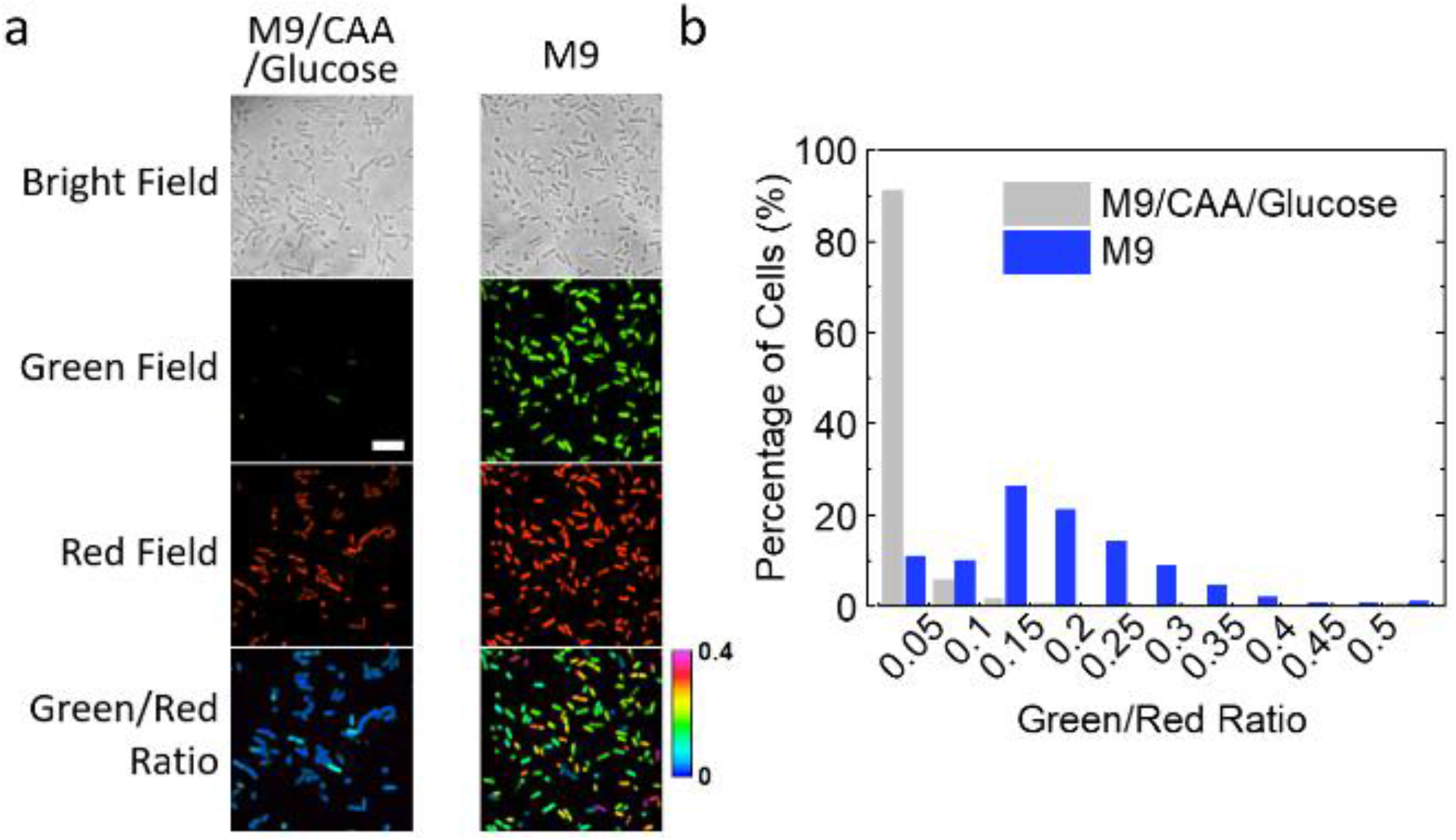
Cellular distributions of (p)ppGpp. **a)** Confocal fluorescence imaging of BL21 Star^™^ (DE3) cells that express both the S2 sensor (green field) and eqFP670 far-red fluorescent protein (red field). These cells were incubated in either M9/0.2% casamino acids/0.4% glucose medium or M9 minimal medium for 90 min before imaging. Pseudo-color ratio images indicated the distributions of green-to-red fluorescence intensity ratios, in the range of zero to 0.4. Scale bar, 10 μm. **b)** Statistical distributions of green-to-red fluorescence intensity ratios as measured from >1,000 cells after growing in either M9/0.2% casamino acids/0.4% glucose medium or M9 minimal medium. Individual cells were binned according to the relative green-to-red fluorescence intensity ratio. The percentage of cells in each bin was plotted.

In summary, we reported here the development of RNA-based fluorescent sensors for the selective and sensitive detection of (p)ppGpp. For the first time, these genetically encoded sensors allowed (p)ppGpp to be directly imaged in living cells, which provided a groundbreaking approach to study the cell-to-cell variations and dynamics of (p)ppGpp biosynthesis and accumulation. Considering the importance of (p)ppGpp in the stringent response of bacteria, we envision that these sensors can be broadly used to study the detailed mechanism of the physiological regulation of bacterial survival during stress, virulence, antibiotic resistance and persistence.

## Acknowledgements

The authors gratefully acknowledge the UMass Amherst start-up grant, NIH R01AI136789, NSF CAREER, and Sloan Research Fellowship to M. You and NIH R35GM130320 to P. Chien. We are grateful to Dr. James Chambers for the assistance in fluorescence imaging and thank Dr. Jade Wang (U Wisconsin) for the gift of the pSM11 plasmid. The authors also thank other members in the You Lab and Chien Lab for useful discussion and valuable comments.

