## Supplementary Information for "Live-Cell Imaging of (p)ppGpp with RNA-based Fluorescent Sensors"

**Table of Contents**

1. Materials and Methods
2. Sequences used for the construction of RNA sensors
3. Sensors design and optimization (S1, S2)
4. Sensor *in vitro* characterization (S3–S8)
5. Live-cell imaging of (p)ppGpp (S9–S12)
6. Sensor plasmid maps (S13)
7. Imaging condition optimization (S14, S15)
8. References

**Materials and Methods:**

**Reagents and equipment.** All the chemicals were purchased from Sigma or Fisher Scientific unless otherwise noticed. ppGpp and pppGpp were purchased from Jena Bioscience (Jena, Germany). DNA oligonucleotides were synthesized and purified by W. M. Keck Oligonucleotide Synthesis Facility (Yale University School of Medicine) and Integrated DNA Technologies (Coralville, IA). All the RNA structures were designed using an Mfold online software(http://www.unafold.org/mfold/applications/rna-folding-form.php). Simulation parameters used are as default. Oligonucleotides were dissolved at 100 μM concentration in 10 mM Tris–HCl, 0.1 mM EDTA at pH= 7.5 and stored at -20°C. Double-stranded DNA template and non-template strands for *in vitro* transcription were prepared by PCR amplification using an Eppendorf Mastercycler. PCR products were gel purified and isolated using a QIAquick PCR purification kit (Qiagen, Germantown, MD). The concentrations of nucleic acids were measured using a NanoDrop One UV-Vis spectrophotometer. RNAs for the *in vitro* experiments were transcribed using a HiScribe™ T7 high yield RNA synthesis kit (New England BioLabs, Ipswich, MA) and then column-purified. All the RNA strands were prepared into aliquots and stored at -20°C for immediate usage or at -80°C for long-term storage.

**Fluorescence assay.** All the *in vitro* fluorescence measurements were conducted with a PTI fluorimeter (Horiba, New Jersey, NJ). Fluorescence assays were conducted in a buffer consisting of 40 mM HEPES, 100 mM KCl, 0.1% DMSO and 5 mM $MgCl_2$, at pH= 7.5. In these measurements, 1 μM RNA and 20 μM DFHBI-1T were used. All the samples were first incubated at room temperature for 90 min before the measurements. The 500–520 nm emission spectra were collected by exciting at 488 nm. Kinetic assays were conducted by exciting at 488 nm and collecting fluorescence signal at 503 nm. All the data were plotted in the Origin software. For fluorescence enhancement and selectivity results (Figure 1b, 1c, S1b, S2, and S3), we normalized all signals to the max value within each figure as 100. For dose-response curves and kinetics results (Figure 1d, 4, and S4–S8), we take the fluorescence before ppGpp addition or after ppGpp removal as the background, and deduct it from all data points within the figure, then normalized to the highest point as 100.

**Vector construction.** S1, S2, and Broccoli (with a T7 promoter and terminator sequence) were cloned, respectively, into a pET28c vector for single-color imaging, or into an eqFP670 fluorescent protein-expressing pETDuet vector for dual-color imaging. The pET28c vector was first digested with BglII and XhoI restriction enzymes (New England BioLabs), and the pETDuet vector was first digested with SgrAI & SacII restriction enzymes (New England BioLabs). After gel purification, the digested vectors were ligated with a similarly digested S1, S2, or Broccoli insert using T4 DNA ligase (New England BioLabs). The ligated products were transformed into BL21 Star™ (DE3) cells (New England BioLabs) and screened based on kanamycin resistance for the pET28c vector or ampicillin resistance for the pETDuet and pBAD vectors. All these plasmids were isolated, and the sequences were confirmed by Sanger sequencing at Genewiz, NJ. For RelA gene validation experiments, both the pSM11 plasmid (IPTG inducible truncated RelA expression; gift from J. Wang) and sensor-expressing pET28c vector were transformed into BL21 Star™ (DE3) cells and screened based on their resistance to both kanamycin and ampicillin. Empty pSM11 vector, pKK223-3, was purchase from Nova lifetech Inc.

**Cellular imaging and data analysis.** Most cellular imaging was performed according to a previously established protocol.[1] BL21 Star™ (DE3) cells that expressing corresponding vectors were grown in LB media at 37°C until $OD_{600}$ reached 0.4 – 0.5, and then 1 mM IPTG was added for a 2 h induction. After induction, the cells were adhered to poly-L-lysine-pretreated glass bottom dish at 37°C for 45 min, and then 200 μM DFHBI-1T was added at room temperature 30 min before imaging. All fluorescence images were collected with a Yokogawa spinning disk confocal on a Nikon Eclipse-TI inverted microscope using an NIS-Elements AR software. Broccoli and RNA sensors were imaged with a 488 nm laser line, while eqFP670 was excited with a 561 nm laser, all through a 60× oil immersion objective.

The RelA strain imaging (Figure 2) was performed on LB-agar pad for both phase-contrast and fluorescence imaging. Here, after 2 h IPTG induction, 200 μM DFHBI-1T was added at room temperature and incubated for 30 min. The cells were then dispensed onto the agar pad. Phase-contrast microscope was used to observe the cell morphology. Both phase-contrast and fluorescence images were collected in a Nikon TiE inverted microscope, which was fitted with an Andor Zyla sCMOS camera. An NIS-Elements AR software was used for the data collection. Cells were excited with a 488 nm laser line through a 60× air objective.

Data analysis was performed with an NIS-Elements AR Analysis software, and the data calculation and fitting were done with the Origin software. Individual cellular fluorescence was analyzed in ImageJ with a MicrobeJ plug-in.

**Table S1.** Sequences used for the construction of RNA sensors. The (p)ppGpp aptamer sequences from *T. oceani 112 ilvE* and *D. hafniense ilvE* riboswitches were shown in black. Broccoli sequences were shown in green. The transducer sequences were bolded and shown in red.

| Sensor | Sequence |
|---|---|
| *T. oceani 112 ilvE* | GGAAGUGUACCUUAGGGUUCCGGCCAUAAGGCGUCAGCGACCGAGCGGUACAAUCCGGGGAAACCCGGAACACCGUGAGCAUAAAAGGCUCCAGCGGCAAGUUCC |
| *D. hafniense ilvE* | GGAAGUGUACCUAGGGUUCCGGGAGCUGCUCCGUCUGGUCCGAGCGGUACAAGAUCCAGAGCAUGGAUUUACACCGUGGGCAGAAAAUACCCGAGCGGAAAGUUCC |
| T1 | GAGACGGUCGGGUCCAGGGAAGUGUACCUUAGGGUUCCGGCCAUAAGGCGUCAGCGACCGAGCGGUACAAUCCGGGGAAACCCGGAACACCGUGAGCAUAAAAGGCUCCAGCGGCAAGUUCCCUGUCGAGUAGAGUGUGGGCUC |
| T2 | GAGACGGUCGGGUCCAG**A**GGAAGUGUACCUUAGGGUUCCGGCCAUAAGGCGUCAGCGACCGAGCGGUACAAUCCGGGGAAACCCGGAACACCGUGAGCAUAAAAGGCUCCAGCGGCAAGUUCC**U**CUGUCGAGUAGAGUGUGGGCUC |
| T3 | GAGACGGUCGGGUCCAGGAAGUGUACCUUAGGGUUCCGGCCAUAAGGCGUCAGCGACCGAGCGGUACAAUCCGGGGAAACCCGGAACACCGUGAGCAUAAAAGGCUCCAGCGGCAAGUUCCUGUCGAGUAGAGUGUGGGCUC |
| S1 | GGAAGUGUACCUUAGGGUUCCGGCCAUAAGGCGUCAGCGACCGAGCGGUACAA**U**CUGUCGAGUAGAGUGUGGGCUCGCAAGAGACGGUCGGGUCCAG**A**ACACCGUGAGCAUAAAAGGCUCCAGCGGCAAGUUCC |
| S2 | GGAAGUGUACCUUAGGGUUCCGGCCAUAAGGCGUCAGCGACCGAGCGGUACAA**UG**CUGUCGAGUAGAGUGUGGGCUCGCAAGAGACGGUCGGGUCCAG**CA**ACACCGUGAGCAUAAAAGGCUCCAGCGGCAAGUUCC |
| S3 | GGAAGUGUACCUUAGGGUUCCGGCCAUAAGGCGUCAGCGACCGAGCGGUACAA**UCG**CUGUCGAGUAGAGUGUGGGCUCGCAAGAGACGGUCGGGUCCAG**CGA**ACACCGUGAGCAUAAAAGGCUCCAGCGGCAAGUUCC |
| S4 | GGAAGUGUACCUUAGGGUUCCGGCCAUAAGGCGUCAGCGACCGAGCGGUACAA**UCGG**CUGUCGAGUAGAGUGUGGGCUCGCAAGAGACGGUCGGGUCCAG**UUGA**ACACCGUGAGCAUAAAAGGCUCCAGCGGCAAGUUCC |
| S5 | GGAAGUGUACCUUAGGGUUCCGGCCAUAAGGCGUCAGCGACCGAGCGGUACAA**UCGG**CUGUCGAGUAGAGUGUGGGCUCGCAAGAGACGGUCGGGUCCAG**UCGA**ACACCGUGAGCAUAAAAGGCUCCAGCGGCAAGUUCC |

| | |
|---|---|
| D3 | GGAAGUGUACCUAGGGUUCCGGGAGCUGCUCCGUCUGGUCCGAGCGGUA CAAUCGCUGUCGAGUAGAGUGUGGGCUCGCAAGAGACGGUCGGGUCCAG CGAUACACCGUGGGCAGAAAAUACCCGAGCGGAAAGUUCC |

These sensors were designed using an Mfold software, which can predict the thermodynamically stable secondary structures of each RNA sequence. Transducers with more base pairs and/or high G/C pair percentage are more thermodynamically stable to induce the proper folding of both Broccoli and the (p)ppGpp-binding aptamer region. The optimum transducer and sensor structure should not be preferably folded in the absence of the target, but in such a near metastable way that could potentially fold with a reasonable small amount of energy, which is assumed to be from the binding of (p)ppGpp. All the tested transducers were designed based on such a rationale, but the real target response still require the experimental testing.

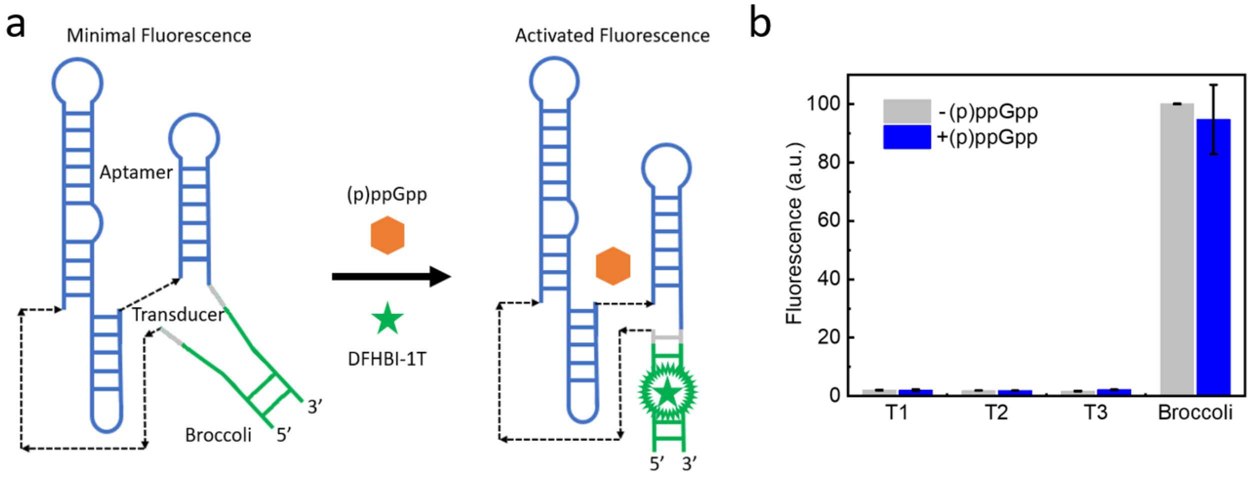


**Figure S1.** Designs of (p)ppGpp sensors. (a) Modular design of the sensors comprises of a Broccoli (green), a (p)ppGpp-binding aptamer (blue), and a transducer sequence (gray). (b) *In vitro* characterization of T1 – T3 sensors. Spectra were measured in a solution containing 1 μM RNA, 20 μM DFHBI-1T, and either 0 or 10 μM ppGpp, at 90 min after mixing. Shown are the mean ± standard deviation of three independent replicates. These results showed that T1 – T3 sensors cannot be used to detect (p)ppGpp.

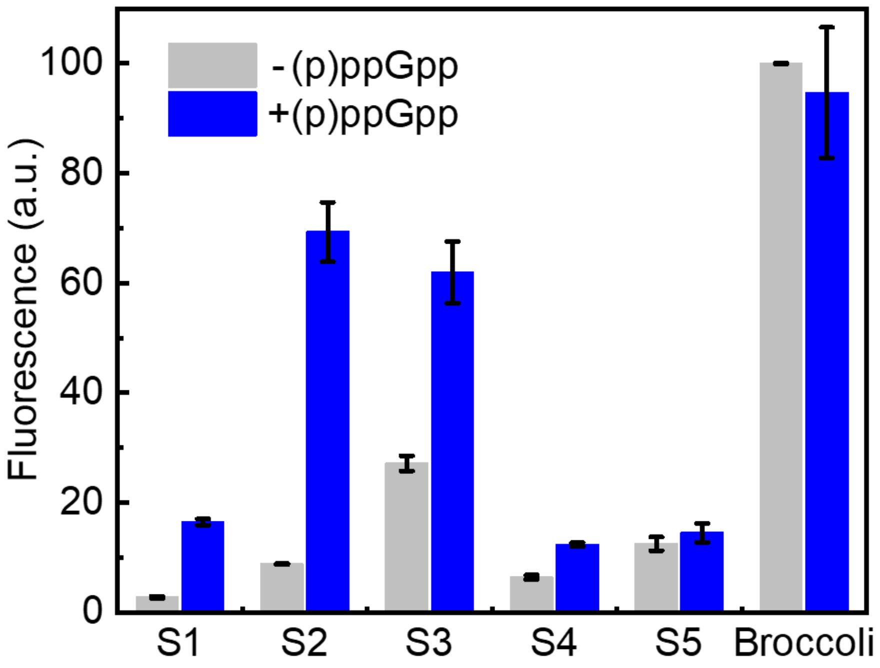


**Figure S2.** *In vitro* characterization of S1 – S5 sensors. Spectra were measured in a solution containing 1 μM RNA, 20 μM DFHBI-1T, and either 0 or 10 μM ppGpp, at 90 min after mixing. Shown are the mean ± standard deviation of three independent replicates. ppGpp-induced significant fluorescence activations were observed with S1 – S3 sensors.

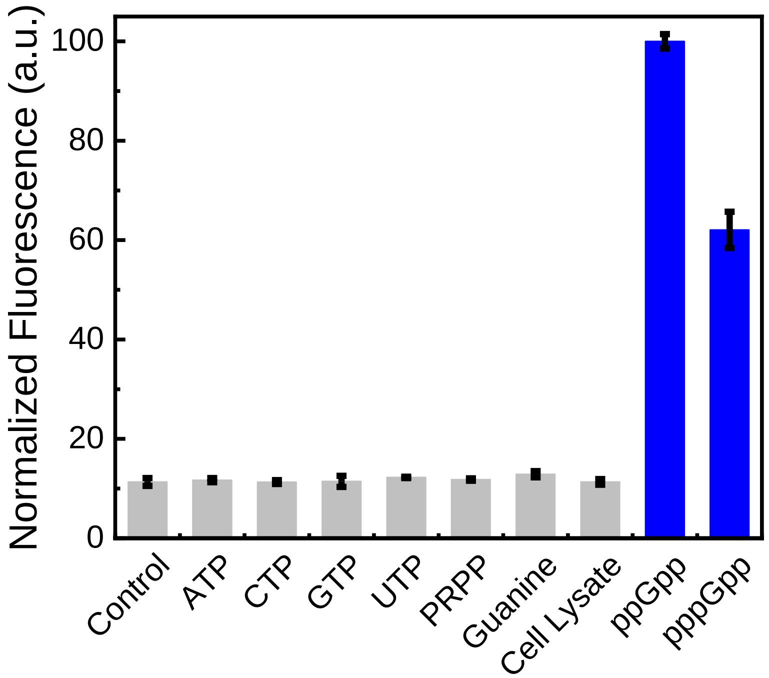


**Figure S3.** Selectivity of the S1 sensor. Spectra were measured with 10 mM NTPs, 10 μM other ligands, or bacteria cell lysate in a solution containing 1 μM S1 and 20 μM DFHBI-1T. Control was measured without adding ligands. Shown are the mean ± standard deviation of three independent replicates.

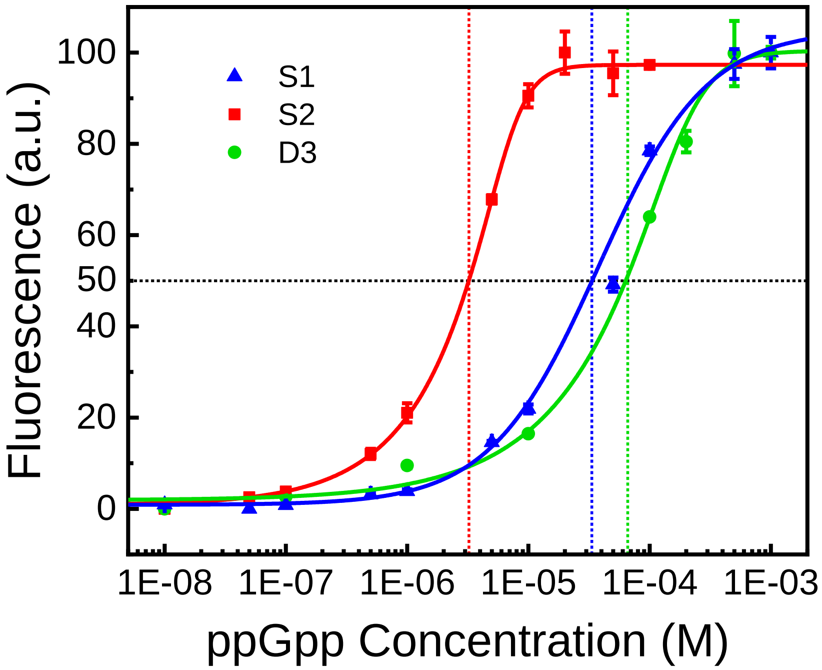


**Figure S4.** ppGpp dose-response curve of the S1, S2 and D3. Spectra were measured in a solution containing 1 µM RNA, 20 µM DFHBI-1T, and different concentrations of ppGpp, at 90 min after mixing. $k_D$ of S1, S2, and D3 are calculated to be 35.6±15.6, 2.7±0.3, and 72.2±14.1 µM, respectively. The limit of detection was calculated to be 0.3 µM. Shown are the mean ± standard deviation of three independent replicates. The difference in detection range between S1 and S2 comes from their different lengths of the transducer. S2 has one more G/C base pair in the transducer region than S1, therefore both the Broccoli and (p)ppGpp-binding aptamer region in S2 are more stable and easier to fold than in S1. The difference between D3 and S sensors comes from their parent riboswitches. D3 adopts a much weaker binding riboswitch sequence than S sensor do, therefore D3 has a general higher detection range than S sensors.

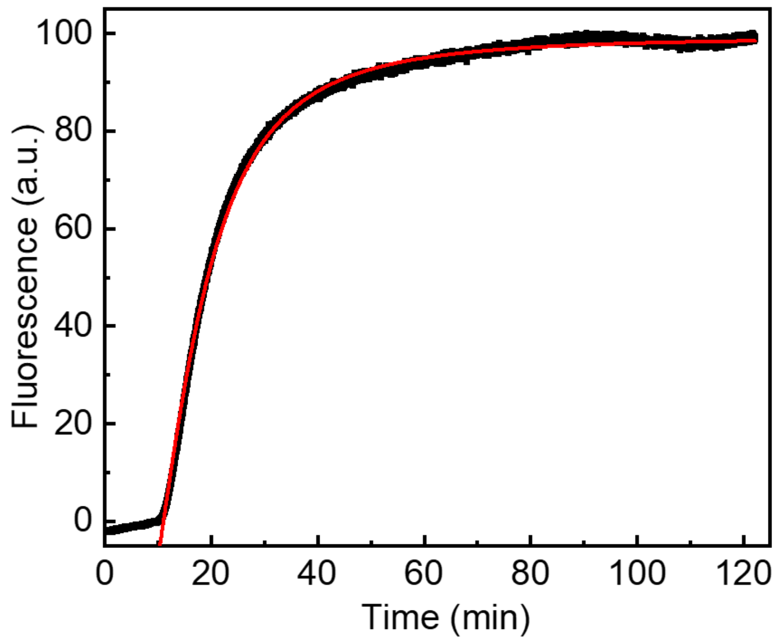


**Figure S5.** Kinetic response of the S2 sensor. At t= 12 min, 10 μM final concentration of ppGpp was added to a solution containing 1 μM S2 and 20 μM DFHBI-1T. The red line indicated the fitted kinetic function. Half-maximum fluorescence level was reached at t= 19 min, and 80% of maximal signal was reached at t= 31 min.

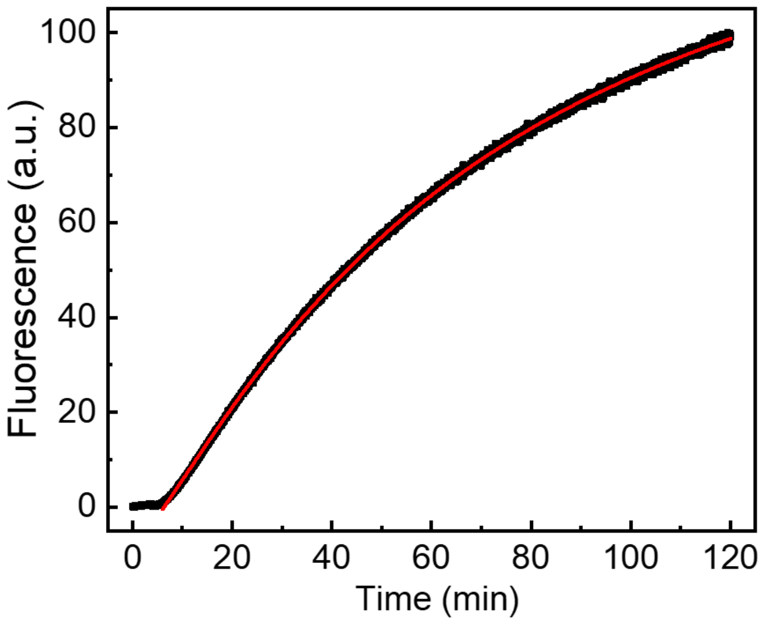


**Figure S6.** Kinetic response of the S1 sensor. At t= 7 min, 10 μM final concentration of ppGpp was added to a solution containing 1 μM S1 and 20 μM DFHBI-1T. Based on the fitted curve function, the maximum fluorescence intensity would be 153. Therefore, the "half-maximum" fluorescence level was reached at t= 72 min, 80% of maximal signal would be reached at t= 212 min, and 90% of maximal signal would be reached at t= 386 min.

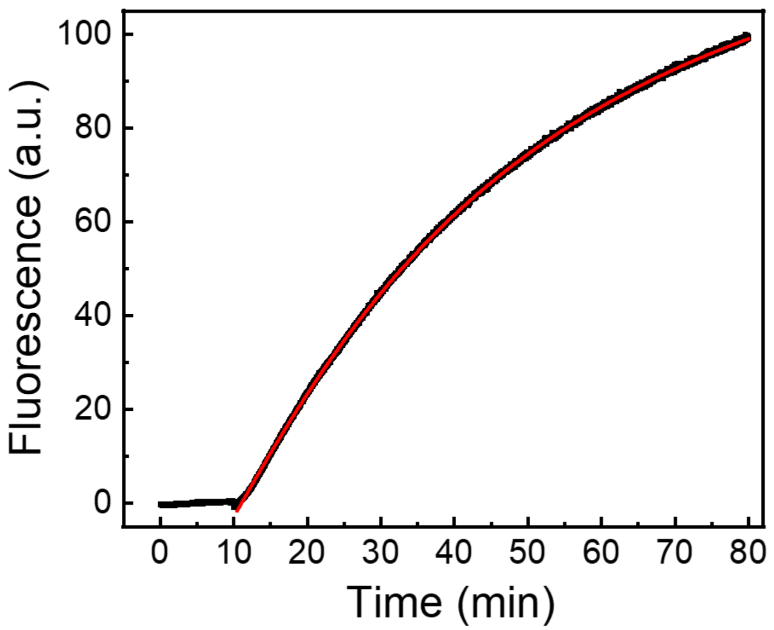


**Figure S7.** Kinetic response of the D3 sensor. At t= 10 min, 10 μM final concentration of ppGpp was added to a solution containing 1 μM D3 and 20 μM DFHBI-1T. Based on the fitted curve function, the maximum fluorescence intensity would be 146. Therefore, the “half-maximum” fluorescence level was reached at t= 49 min, 80% of maximal signal would be reached at t= 125 min, and 90% of maximal signal would be reached at t= 223 min.

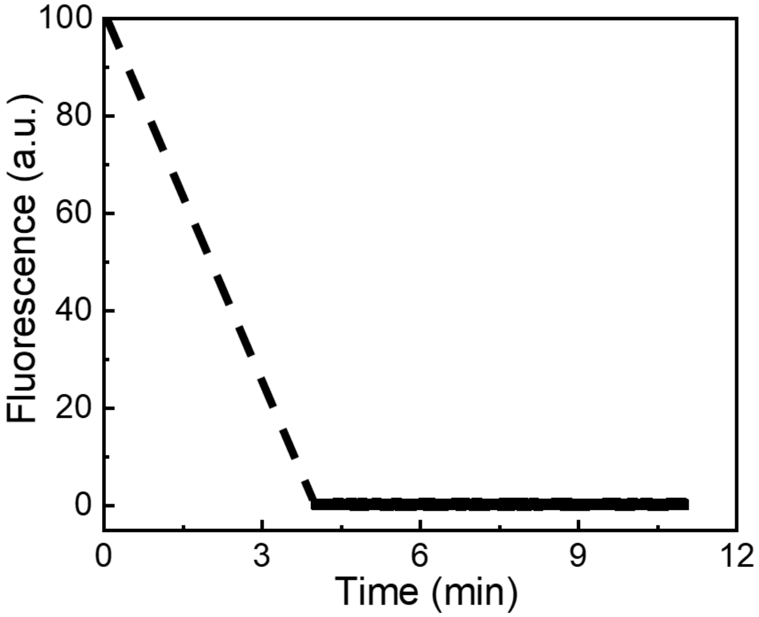


**Figure S8.** Fluorescence deactivation rate of the S2 sensor. After mixing 10 μM ppGpp with 1 μM S2 and 20 μM DFHBI-1T for 1 h, the solution was centrifuged through a desalting column (Bio-Rad Laboratories, Inc) to remove free ppGpp in approximately 4 min. This desalting column has been pre-treated to contain 40 mM HEPES, 100 mM KCl, 0.1% DMSO, 5 mM $MgCl_2$, and 20 μM DFHBI-1T, at pH= 7.5, the same solution as that of the original S2/ppGpp mixture. These results indicated that the fluorescence deactivation took less than 4 min after removing free ppGpp.

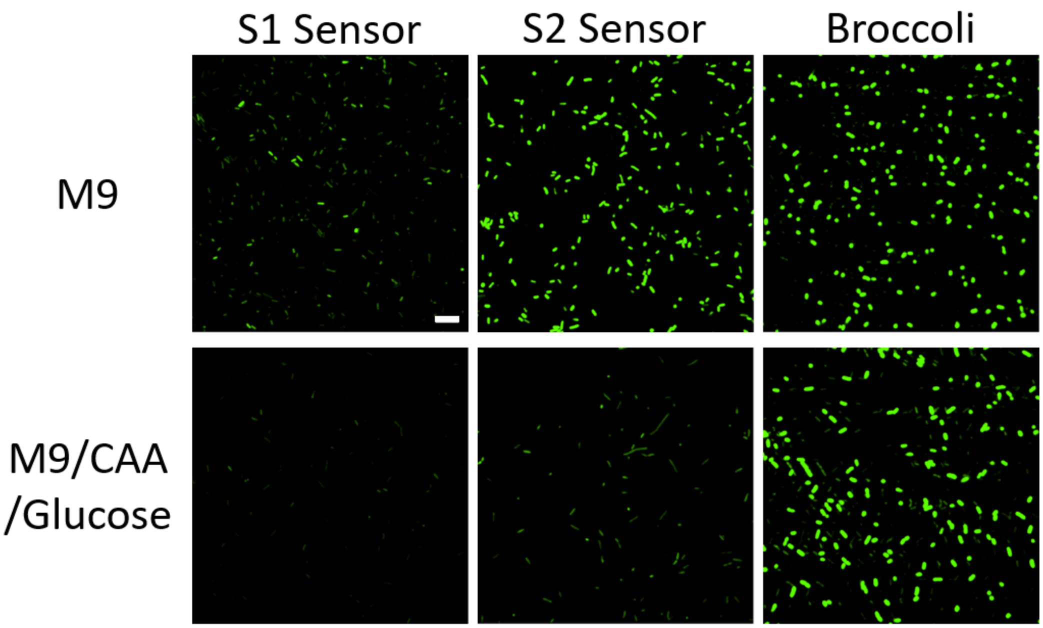


**Figure S9.** Imaging of RNA sensor- or Broccoli-expressing BL21 Star™ (DE3) cells under different nutritional conditions. Images were taken 1 h after incubating with 200 µM DFHBI-1T in either M9 minimal medium (limited nutrients and high (p)ppGpp level) or M9 medium supplemented with 0.2% casamino acids (CAA) and 0.4% glucose (rich nutrients and low (p)ppGpp level). As expected, S2 fluorescence was in general higher than that of S1. Broccoli signal functioned as a control and exhibited no obvious difference under these two nutritional conditions. Scale bar, 10 µm.

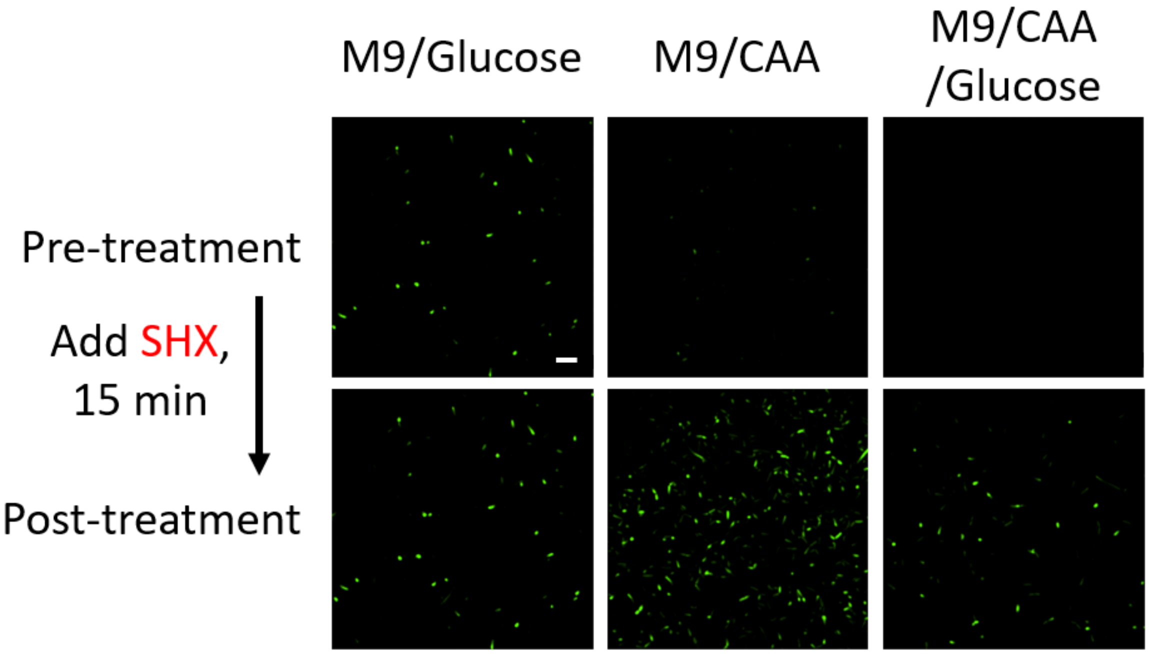


**Figure S10.** Serine hydroxamate (SHX)-induced (p)ppGpp biosynthesis. SHX functions by inducing the starvation of amino acids. S2-expressing BL21 Star™ (DE3) cells were first incubated in M9 medium supplemented with 0.4% glucose, 0.2% casamino acids (CAA), or both nutrients. Afterwards, 1% final concentration of SHX was added 15 min before imaging. Scale bar, 10 µm.

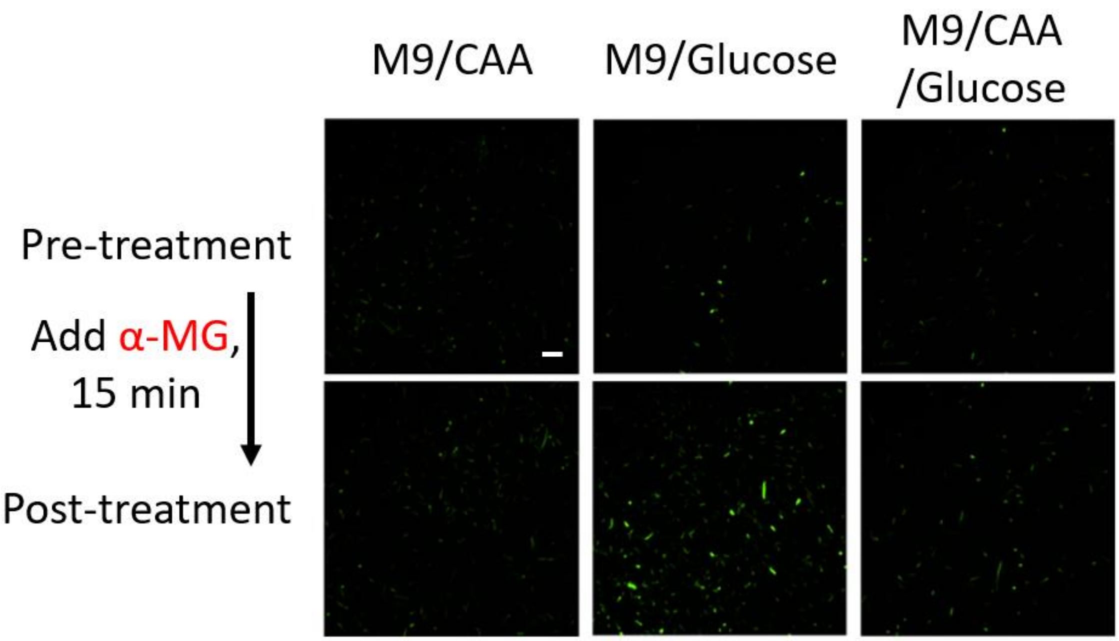


**Figure S11.** Methyl-α-glucose (α-MG)-induced (p)ppGpp biosynthesis. Here, α-MG functions by inducing the starvation of glucose. S2-expressing BL21 Star™ (DE3) cells were first incubated in M9 medium supplemented with 0.4% glucose, 0.2% casamino acids (CAA), or both nutrients. Afterwards, 2.5% final concentration of α-MG was added 15 min before imaging. Scale bar, 10 μm.

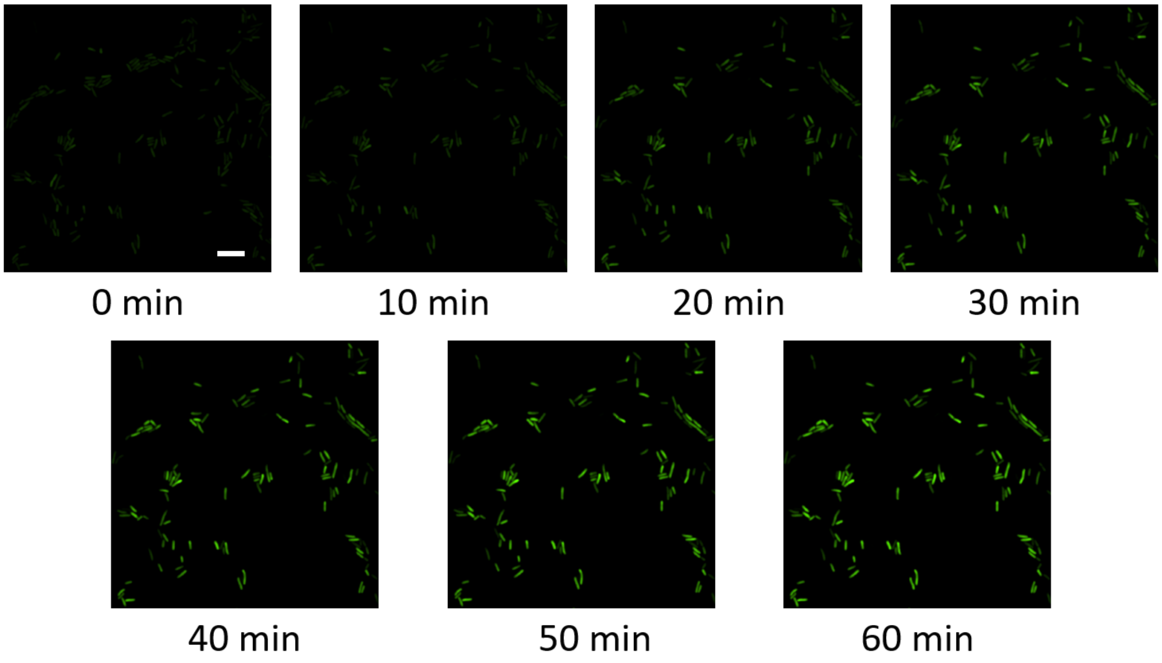


**Figure S12.** Kinetics of (p)ppGpp biosynthesis in *E. coli* cells. Confocal fluorescence imaging of S2-expressing BL21 Star™ (DE3) cells after switching at 0 min from M9/0.2% casamino acids/0.4% glucose medium to M9 minimal medium. Scale bar, 10 μm.

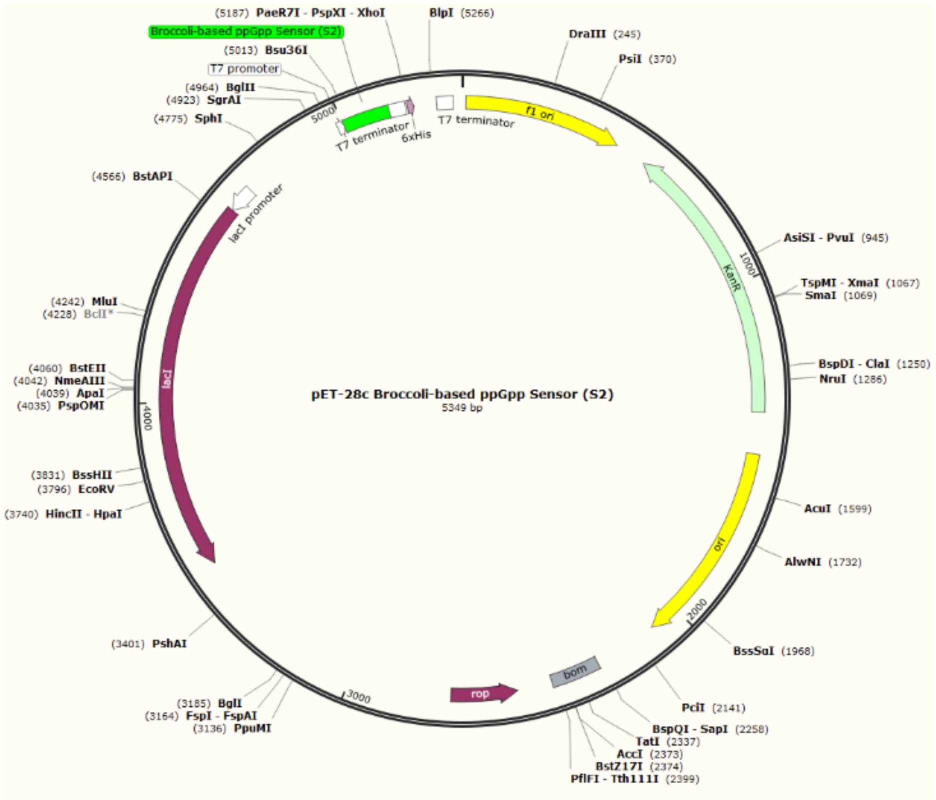


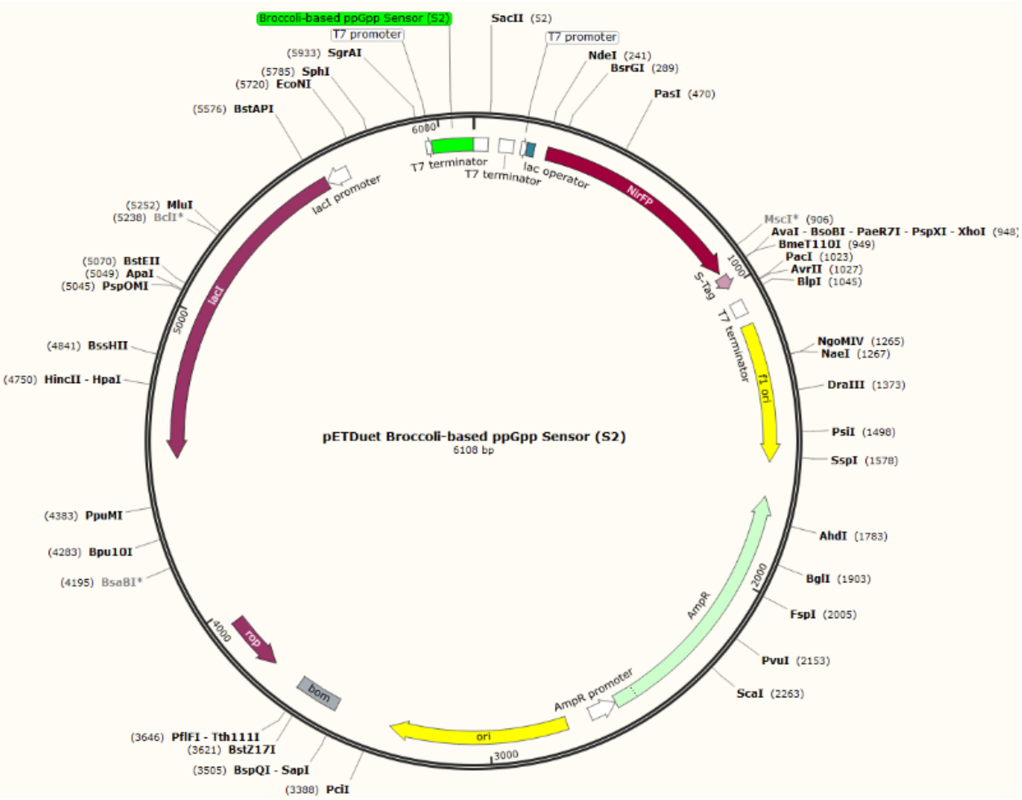


**Figure S13.** Plasmid maps of the S2 sensors. Top, a pET28c-based S2 sensor plasmid map. Bottom, a pETDuet-based plasmid that express both S2 sensor and eqFP670, a far-red fluorescent protein (NirFP). Both sensor and NirFP expressions are under the control of p(T7) and induced by adding IPTG.

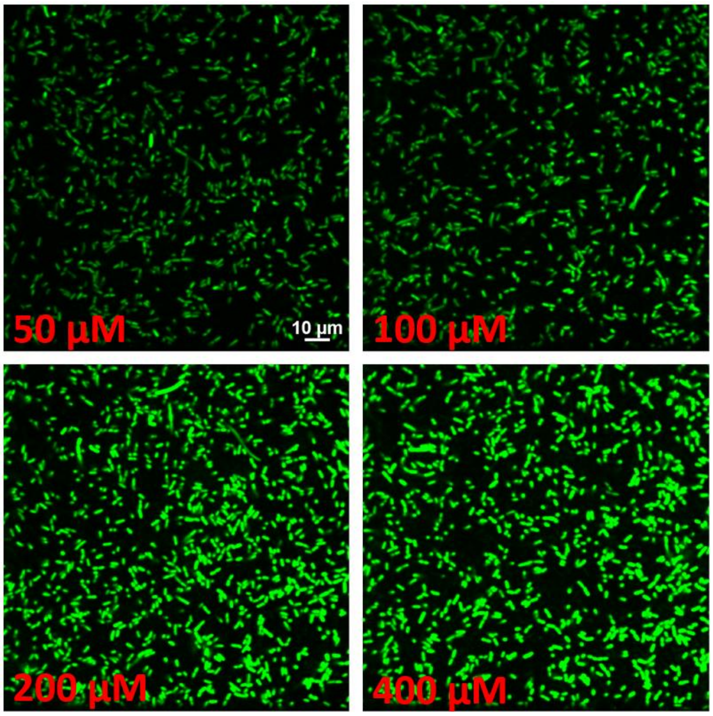


**Figure S14.** The choice of DFHBI-1T concentration in cellular imaging. Broccoli-expressing BL21 Star$^{TM}$ (DE3) cells were imaged after adding 50–400 µM of DFHBI-1T. We chose to use 200 µM concentration of DFHBI-1T for cellular imaging in this study considering its bright but not oversaturating fluorescence signal. Scale bar, 10 µm.

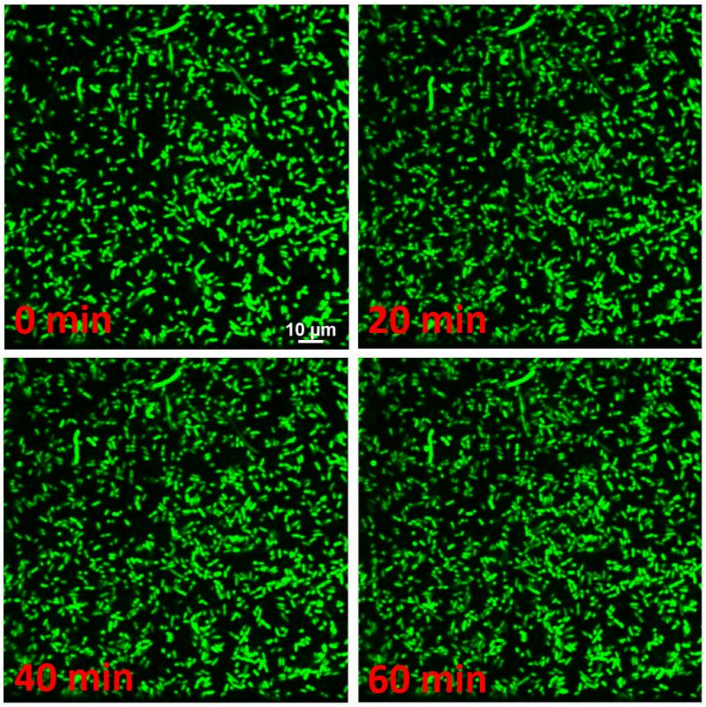


**Figure S15.** Photobleaching of cellular fluorogenic RNA signals. Broccoli-expressing BL21 Star$^{TM}$ (DE3) cells were imaged at 5 min interval over a total of 60 min period. 0, 20, 40, and 60 min images were shown as representative. No obvious photobleaching effects were shown. Scale bar, 10 µm.
